# Benchmarking the robustness of segmentation models to corruptions in biological imaging

**DOI:** 10.64898/2026.08.21.746302

**Authors:** Yekta Kesenci, Loïc Le Folgoc, Elsa Angelini

## Abstract

Deep-learning-based segmentation algorithms have gained considerable accuracy for processing biological images. In particular, the introduction of large foundation models, novel architectures, and semantically varied datasets now allows for deployment of state-of-the-art models for clean image cohorts with limited re-training or, in the best of cases, in an out-of-the-box fashion. Biological imaging, however, is liable to corruptions that can hinder their deployment. While some methods document their robustness to the most common corruptions, a systematic robustness analysis of the state of the art to the expansive gamut of corruptions in biological imaging remains to be done. We perform this benchmarking by simulating 36 corruption types with varying degradation severity on images sampled from 30 different datasets. Our benchmark accounts both for the variety in biological images and the nature of corruptions. Among other things, our study reveals that performance on clean images does not correlate with overall robustness to image corruptions. In fact, we find that a decade-old method, StarDist, is more robust than many of its more recent foundation-model-based counterparts. We also show in a dedicated representation analysis that the performance of segmentation models collapses in the early layers of the encoding phase.

## Introduction

Cell segmentation is the preliminary step of many image processing protocols, such as morphology analysis (1), cell tracking, or mechanobiology (2). It is also a difficult task that admits no universal solution to the varying experimental conditions and cell types that biologists face (3). Deep-learning-based methods have made progress towards that goal, thanks in part to the availability of large annotated datasets (4), the introduction of architectures specifically designed for the segmentation of cellular structures (5), and the use of large foundation models with strong inductive biases (6). However, to be considered deployable, these methods ought to be robust against optical corruptions occurring in biological imaging.

Every step of the imaging process, from the design of the experimental setup to the digital recording of the acquisition, is susceptible to corruptions (7), (8). Annotated datasets include some corrupted images, but in unbalanced proportions that only partially reflect real experimental variety. Although some segmentation methods report their robustness against the most common types of corruptions (9), no systematic evaluation of the robustness of the state of the art has been published — except for (10), on a limited number of corruptions and adversarial attacks, and for simulated images and fluorescence images only.

Benchmark studies proved nevertheless valuable in related deep-learning tasks. The seminal study of (11) showed that the robustness of classifier models to common camera perturbations is not correlated with clean image performance and advocated the use of multiscale architectures. In contrast, (12) showed that such correlation existed with semantic segmentation models. Self-supervised learning has been shown to moderately increase segmentation performance in clean images, but (13) demonstrated that it greatly increased classification robustness. We can see that these studies yield different conclusions for different downstream tasks. Cell segmentation, on the other hand, is quite different from any of these tasks: it aims to distinguish an arbitrarily high number of similar cells in a variety of image types. The robustness of cell segmentation methods cannot therefore be extrapolated from existing studies.

We propose a benchmarking protocol to assess the robustness of several state-of-the-art segmentation algorithms in biological imaging. Starting from a curated pool of 500 images sampled to maximize diversity from 30 public datasets spanning 7 imaging modalities, 12 staining protocols, and 27 cell types, we construct a corrupted dataset of 90 500 images by simulating 36 corruption types, grouped into five physically-motivated categories (noise, blur, geometric distortion, digital compression artifacts, and assay-specific artifacts), at five severity levels each. We use this corrupted dataset to evaluate the global performance of several state-of-the-art segmentation algorithms, and design specific metrics to assess their robustness against the simulated corruptions. Our study shows that the segmentation models that perform best on clean images with varied semantic properties are not necessarily the most robust. For example, we show that StarDist (14) is significantly more robust than some of its more recent SAM-based counterparts. To explain these differences, we conduct a representation layer comparison of the Cellpose and StarDist architectures and show that segmentation performance collapses in the early layers of their networks’ architectures.

## Results

### Design of the corrupted dataset

We start by creating a “corrupted” dataset of biological images with their ground truth segmentation masks. To be a reliable yardstick of deployability, it shall feature both images with semantically varied information and corruptions of gradual severity. We proceed in two steps.

#### Image sampling

First, we sample 500 biological images with optimized diversity (see Figure 1a). We choose them from 30 public datasets that account for 7 imaging modalities, 12 staining protocols, and 27 cell types. Collecting these datasets guaranties inter-dataset variability. To ensure that the chosen images are as diverse as possible within these datasets (intra-dataset variability), we deploy a two-step farthest point sampling strategy (15). All images are first embedded with the small pre-trained Dinov2 transformer (distilled ViT-S 14) (16). We then select 10 images from each dataset using a greedy algorithm that maximizes the orthogonality of their representations. Our second step repeats this process, but this time by sampling from the total number of images contained in all datasets. We refer to the Methods section for further elaborations on the selected datasets and the sampling strategy. Our framework not only collects images with diverse semantic qualities (see Figure 1c), but also ensures that these images present varying segmentation challenges: Figure 1b shows that their ground-truth segmentation masks can feature both high as well as low numbers of ROIs, foreground to background ratio, a nearest-neighbor distances between ROIs, and a spatially unbiased ROI distribution.

**Fig. 1.**
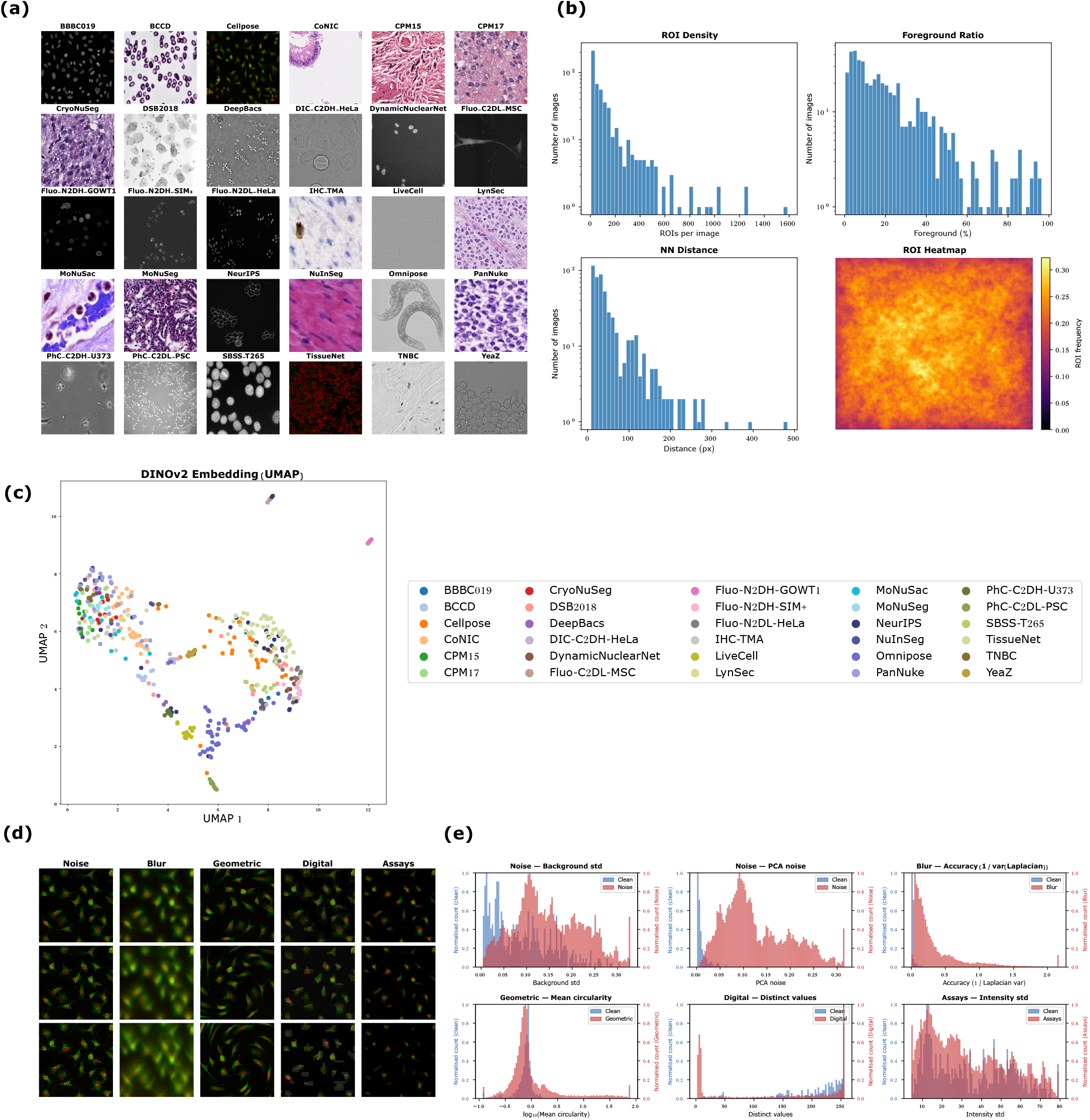
**a**, Sample crops of each considered datasets. **b**, Statistics over the masks of the gathered clean images: the number of ROIs per image, the ratio of foreground (union of all ROI pixels) to background in percentage, the mean nearest-neighbor distance of each ROI mask per image, the probability of each pixel of an image to be an ROI. **c**, The first two components of the UMAP algorithm applied on the embedding of the DINOv2 ViT-S/14 pre-trained model for each image of the clean dataset (each grayscale image was concatenated three times to create three channels). **d**, Examples one sample per corruption category, applied on a single image from the Cellpose dataset with increasing severity: speckle (Noise), coma (Blur), shear (Geometric), JPEG compression (Digital), photobleaching (Assays). **e**, Corruption statistics over each image of the clean and corrupted dataset: from left to right, top to bottom, standard deviation of the background, mean of the last ten principal components of the image, accuracy of the image (inverse of the variance of the Laplacian of the image), mean circularity (4*π*Area*/*Perimeter^2^) of each ROI, the number of distinct values within each image, standard deviation of the image.

#### Corruption simulation

The collected 500 images constitutes our “clean” dataset. Although these images naturally contain corruptions, they are not representative of both the variety and the severity of the possible corruptions that occur in microscopy imaging. We therefore extend this “clean” dataset into a “corrupted” one by simulating 36 corruption types with 5 severity levels for each. These corruptions can be grouped into 5 categories. *Noise*: stochastic-type of corruptions that occur before the analog conversion of the image. They arise from natural random fluctuations caused by thermal variations in cameras, low-light conditions, short exposure times, or interference of coherent light. *Blur*: blurs arise from the wrong convergence of the optical paths of the illuminated specimen unto the focal plane. Although infinitely varied in practice, they were subjected to a systematic classification by Zernike, from which we include the most common modes, along with the standard Gaussian blur. *Geometric*: cell shape distortions induced by a misalignment of the imaged sample with the optical apparatus. We consider both linear and nonlinear geometric transforms to simulate ROI shapes not seen during training. *Digital*: happens after the analog conversion and includes compression artifact (e.g. JPEG) due to storage limitations and image transfer constraints common in biology. *Assays*: corruptions directly related to the experimental conditions of the biological samples. We define 5 severity levels per corruption types (see Figure 1d), controlled by a single parameter of their mathematical model. Parameter values were set empirically. The lowest severity level is hardly noticeable by a user, while the highest level exceeds what would normally be mandated by scientific standards of imaging acquisition. In all cases, we make sure that the images can be accurately segmented by the human eye. We refer to the Methods section for a comprehensive list of all corruption types, the used mathematical models, and the values of the severity level parameters. Our corrupted dataset includes 90,500 images, and features a considerably wider range of corruption levels as measured by standard noise-estimating metrics compared to the clean dataset (see Figure 1e and Supplementary Figures S4-S8).

### Overall performance of segmentation algorithms

We use our corrupted dataset to evaluate 10 pre-trained deep-learning segmentation models: Cellpose-SAM (9), Cellpose (the ‘cyto3’ model) (17), CellSAM (18), micro-SAM with its 3 transformer sizes variants (ViT-T, ViT-B, ViT-L) (19), StarDist (14), and the baseline SAM model with 3 transformer size variants (ViT-T, ViT-B, ViT-H) (20). These segmentation models have been trained on different datasets, some of which overlap with our own (see Extended Table 2). We denote by “Clean” the subset of the clean images of our dataset, by “All” the entirety of our dataset (with both clean and corrupted versions of each image), and by “Train” the respective subsets of our dataset that includes only the images (clean and corrupted) coming from datasets on which the corresponding segmentation algorithm was trained on.

We first measure segmentation performance over the clean and corrupted datasets using the F1 score (see Methods), a detection metric that accounts for the precision of the produced masks of each ROI. Unless indicated otherwise, the F1 scores are computed with an IoU threshold of 0.5. Results (Figure 2a) confirm several statements that have been previously established in the recent literature: SAM-based models perform better than their non-foundation-model-based counterparts; using larger transformers slightly improves micro-SAM’s performance; on its own, SAM-auto performs poorly, and specialization on biological images is very beneficial; remarkably, the non-transformer-based Cellpose model performs better than all other models, except for Cellpose-SAM and Micro-SAM-L; there is a noticeable model performance difference between clean and corrupted images; all models generalize well to datasets not seen during training, except for StarDist which has been trained on a very limited number of datasets. Based on these results, we can distinguish three performance regimes: the best performing tier, uniquely composed of Cellpose-SAM (F1 > 0.4 on “All”); the second tier, composed of micro-SAM-L, Cellpose, micros-SAM-B (F1 > 0.3); and the third tier, which includes the rest. In a word, given a biological image of arbitrary noise level and type, the Cellpose-SAM model offers the best chance of segmenting it faithfully, at the present date.

**Fig. 2.**
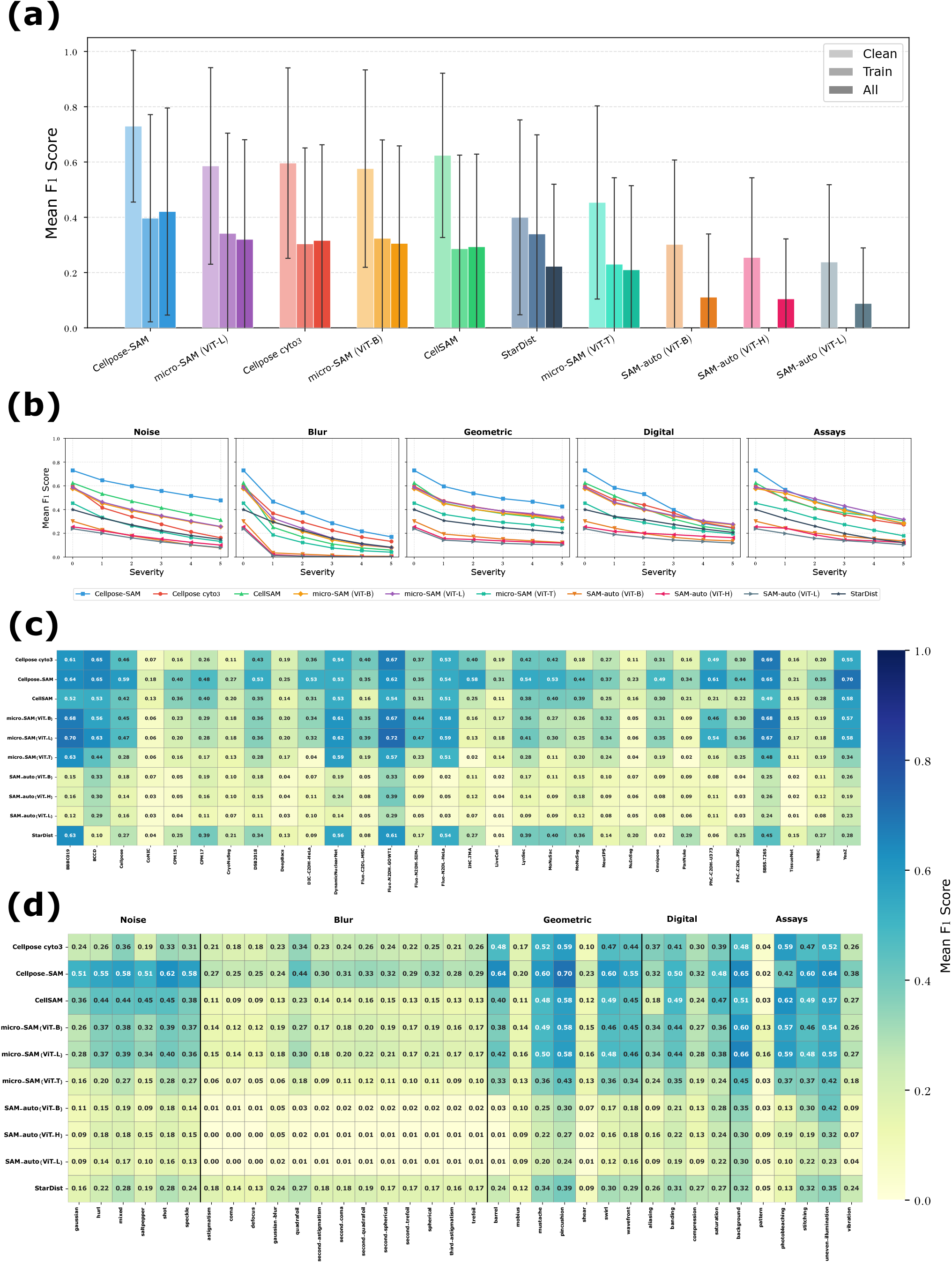
**a**, Mean F1@50 score over the clean dataset, corrupted dataset and the subsets of corrupted training images of each considered segmentation model; models are ordered from best-performing to worst performing over the entirety of our dataset (“All”). **b**, Evolution of the mean F1@50 score, taken over the image and corruption types, against increasing corruption severities for each model and each corruption category. **c**, Mean F1@50 score of each segmentation model for each origin dataset over all corruption types, all corruption severities and all images of the corresponding dataset. **d**, Mean F1@50 score of each segmentation model for each corruption over all images and all severities.

Results in Figure 2b report the degradation of the F1 score against severity levels per corruption type. Overall, at low severity levels, Cellpose-SAM remains the best performing and the 2nd tier group is mostly preserved across severity levels. The differences in model performance are greatly reduced at high severity level, StarDist becoming even competitive with high levels of blur and digital corruptions. Performance degradation patterns reveal 3 types of corruption sensitivity: high corruption sensitivity for Blur, moderate for Digital and Assay corruptions, and low for Noise and Geometric corruptions.

Figure 2c shows the mean F1 scores per dataset over all corruption categories and severity levels per image sub-datasets. Among the best and 2nd tier models, we observe striking differences in F1 performance across datasets. We can draw three groups of them: poorly handled (F1<0.4) with all models – CoNIC, CryoNuSeg, LiveCell, TissueNet, TNBC; strongly handled (F1>=0.6) with at least one model – BBBC019, BCCD, Cellpose, Dynamic Nuclear Net, Fluo-N2DH, Fluo-N2DL, IHC-TMA, SBSS-T265, YeaZ; and others. Looking at the performance of each model, the Segment Anything variants prove uniformly poor over all datasets. This confirms our previous insights, namely that the semantic properties of biological images are fairly distinct from those used for the training of SAM, and that specialized models and training are mandatory. StarDist again emerges as competitive (F1 >= 0.6) for some sub-datasets (BBBC019, Dynamic Nuclear Net, Fluo-N2DL). Also, when we consider datasets where most models perform relatively well (for example BBBC019, DynamicNuclearNet, Fluo-N2DH-GOWT1), we can see that their performance do not differ as significantly as before. Instead, if Cellpose-SAM and micro-SAM-ViT-L do better than the other models overall, it is mostly because their performance is maintained on a greater variety of datasets. This confirms the recent progress in these new models in the recent literature: their performance gains are largely explained by their ability to generalize to new image cohorts.

Finally, Figure 2d reports the average F1 scores over all severity levels per corruption subtype within the 5 corruption categories. Regarding best and 2nd tier models on clean images: (low corruption sensitivity) Noise sub-types degrade similarly all models. Two geometric corruptions (Möbius and Shear) affect drastically segmentation performance for all models; (moderate sensitivity) Pattern and Vibration subtypes of assay corruption affect drastically segmentation performance for all models, while Aliasing and Compression subtypes of digital corruption affect more strongly CellSAM; (high corruption sensitivity) Quadrafoil blur subtype is better handled than others by all models, and StarDist joins the 2nd tier performance level; the non-linear Möbius transform is harder than its linear counterparts; and pattern corruptions are significantly harder compared to other assay-specific corruptions.

### Robustness of segmentation algorithms

We adapt (see Methods) two performance robustness metrics introduced first in (11), and extended for semantic segmentation in (12), for our use-case on biological image segmentation: corruption degradation (CD) and relative corruption degradation (rCD). Both metrics aim to make segmentation models mutually comparable by measuring their robustness to corruption degradation with respect to a reference model, which we have chosen to be StarDist. For a given model and corruption type, CD measures the ratio of the averaged detection errors (be it False Positives or False Negatives) at all severity levels, over the same average obtained with the reference method. This metric also includes errors obtained on clean images. For a given model and corruption type, rCD measures the relative amount of decline of performance as the ratio of the shift of detection errors between corrupted and clean images, over again the same shift for the reference model. CD and rCD are always equal to 1 for StarDist by definition. Values above 1 indicate a lower robustness relative to StarDist, and below 1 a greater robustness relative to StarDist.

Figure 3a reports the CD and rCD values versus the mean F1 score on clean images for each segmentation model and per corruption category. CD is overall correlated to the mean F1 score on clean images (i.e. models preserve their rank), although more strongly so for noise and geometric corruptions (previously identified as low corruption sensitivity) than blur, digital and assays. This confirms our previous intuition for these two corruption types, as documented by the performance discrepancy outlined in Figure 2b. Cellpose appears more affected than other 2nd-tier models for Noise corruptions. rCD, on the other hand, is much more decorrelated to the F1 score on clean images. Focusing on the best and 2nd-tier models, rCD values are above 1 for all cases, except micro-SAM (L and B) for the Assay corruption. For Digital corruptions, we see a clear correlation between the baseline F1 score and the amount of robustness decline: the higher the baseline F1 score the worst the robustness to corruption. For the other corruption types, the models showing best and worst robustness vary. But we note that Cellpose-SAM (which included noise augmentations during its training) is more robust than others for noise corruption (close to StarDist robustness). But Cellpose-SAM also included blur and aliasing augmentations that did not translate in robustness gains versus StarDist. Finally, micro-SAM-T (lower tier) shows both robustness metrics close to StarDist for the Assay corruption type.

**Fig. 3.**
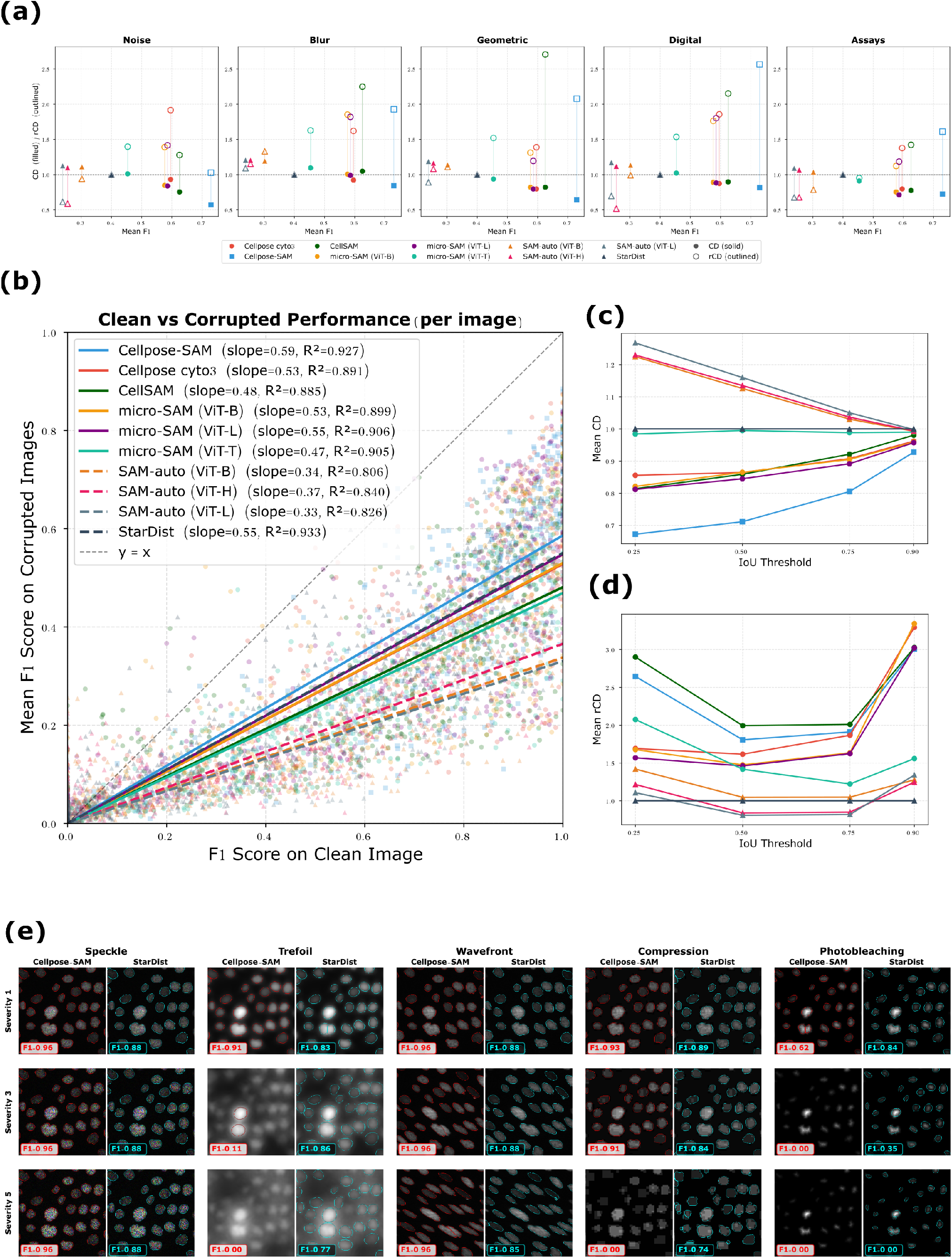
Robustness of the different segmentation models over the corrupted datasets. **a**, CD and rCD of each segmentation model for each corruption category. **b**, F1@50 score of each segmentation model for each clean image against the mean F1@50 score of the respective corrupted versions of each clean image. The slope and *R*^2^ values of the linear regressions with no biases of each model is provided. Third-tier models are dashed. **c-d**, Evolution of the mean CD and mean rCD against the IoU threshold (legend is the same as in Figure 3a); for the rCD metric at IoU = 0.25, we removed images featuring Pincushion deformation as StarDist showcased near-perfect robustness, creating outliers. **e**, Qualitative examples illustrating the decoupling between clean performance and corruption robustness: for five corruption types at severity levels 1, 3, and 5, Cellpose-SAM (left column of each pair) and StarDist (right column) predictions are shown with F1@50 scores overlaid.

We next examine potential biases in the robustness metrics that we used. We first check whether they favor methods that perform poorly on clean images. Indeed, we can suppose that methods that perform very well on clean images have more “room to fall”. In other words, a method that scores 1000 True Positives will lose more True Positives with corruption degradations than another method that scores 400. We empirically check this in Figure 3b. It assigns for each segmentation method 500 pairs of scores: the F1 score on each clean image along with the mean F1 score on the degraded variants of each of these images. A perfectly robust model will have a slope equal to 1: it performs as well on clean images and on their degraded counterparts. This metric partly confirms what was previously established: the StarDist (3rd-tier) model is largely more robust than all second-tier and 3rd-tier models. Cellpose-SAM (1st-tier), on the other hand, appears this time more robust than StarDist. We also document the bias of our metrics to our choice of the IoU threshold for the F1 score. In Figure 3c, we observe that all CD values converge to 1 for high IoU threshold values (0.9), reaching a robustness similar to that of StarDist. This is because their overall performances collapse across images and noise types, making their robustness indistinguishable. When decreasing the IoU threshold, more tolerance is allowed on pixel-level segmentation errors, with the potential to decrease FN (missed detection) but also increase FP. For best performing methods (CDs < 1), Cellpose-SAM is best, followed by the 2nd-tier methods and CellSAM. In Figure 3d, the difference in robustness between StarDist and all best performing models (first-tier and second-tier) is confirmed and amplified. The relative greater robustness of the SAM models is called in question for lower and higher values of the IoU threshold.

We illustrate in Figure 3e visual examples of segmentation results from the best performing model (Cellpose-SAM) and our reference model for robustness (StarDist) on a fluorescence microscopy image with different corruption types and severity levels. We see that, although Cellpose-SAM is considerably better than StarDist in clean or low-level degradation conditions, the quality of its segmentations collapses rapidly with corruption severity, even when images present no challenges for the human eye, whereas StarDist is on par with its estimation on clean image.

Overall, StarDist appears more robust than most (if not all) of its counterparts. Yet, it was not trained on a more extensive training dataset nor leveraged a large foundation model: these strategies, useful for clean images, do not appear profitable to increase model robustness. On the other hand, StarDist uses a strong prior on cell shape, as star-like convex polygons which we posit provides greater robustness properties.

### Intermediate layer analysis

We conclude our benchmarking with an analysis of the robustness of the layers within a segmentation model’s network architecture. To do so, we rely on the Centered Kernel Alignment (CKA) metric (21) which measures how much a pair of layers agree in assigning the same encoding similarity to a pair of image inputs (see Methods). CKA requires a set of images to be computed. We chose the Cellpose dataset (540 images) for its semantic variability. We will also leverage corrupted variants of it. To preserve the same number of images (and the underlying semantic distribution of the Cellpose dataset), we will select a single representative corruption type at severity level 5 for each corruption category: Hurl corruption for Noise, Gaussian blur for Blur, Wavefront distortion for Geometric, JPEG compression for Digital, and Photobleaching for Assays.

In Figure 4, we report CKA-based results comparing Cellpose, the best performing model that does not rely on SAM, with StarDist, which is less accurate on clean images yet more robust. From thereon, we will consider layers that are the output of a 2D convolution, a batch normalization, or a ReLU.

**Fig. 4.**
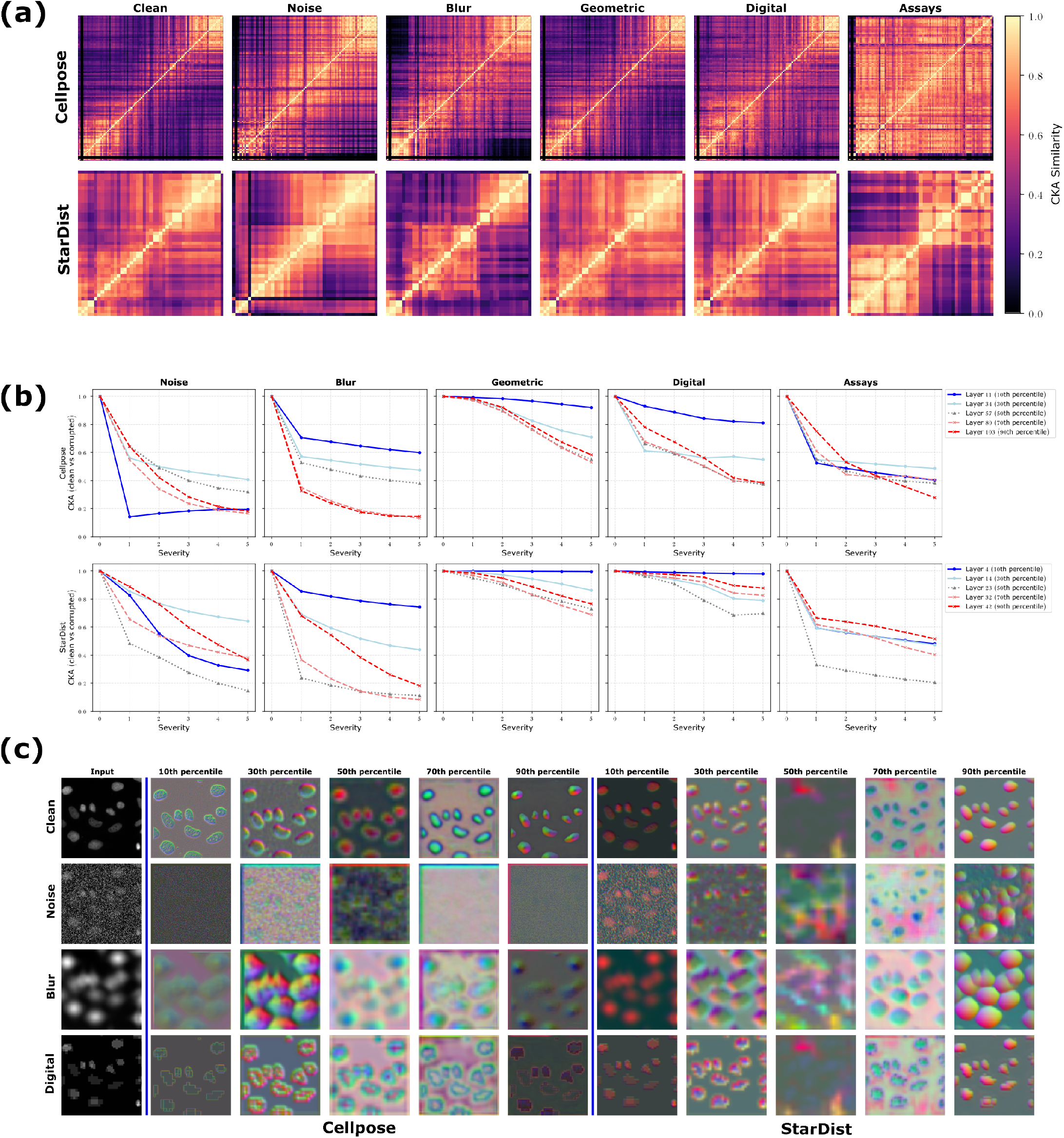
Layer-level effects of image corruptions for the Cellpose and StarDist segmentation models. CKA similarity metric is reported for inter and intra-layers correlations at 3 levels (corresponding to different percentile depths): Encoding path (10th & 30th), Bottleneck (50th) and Decoding path (70th & 90th).(a) Inter-layer CKA similarity computed over the clean dataset and its corrupted variant per corruption type at severity level 5; (b) Intra-layer CKA similarity computed over the clean dataset and its corrupted variant per corruption type and severity level; (c) Feature maps produced by 5 layers of different depth on a clean sample image (taken from BBBC019) and its corrupted counterparts at severity level 5. For each corruption category, a single corruption type is chosen in panels (a) and (b): Noise=Gaussian, Blur=Coma, Geometric=Wavefront, Digital=Compression, Assays = Photobleaching. For panel (c), PCA for RGB encoding and resizing were applied on each feature map.

Figure 4a reports inter-layer similarity matrices on the clean dataset and its corrupted counterparts. Cellpose, whose network is a modified Residual U-net, contains 116 layers, while StarDist, based on a standard U-net, contains 47 layers.

On clean images, Cellpose and StarDist feature common inter-layer similarity structures: shallow layers (those at the beginning of the encoding stage and those at the end of the decoding stage) are alike, while deeper layers bear a greater level of dissemblance to the rest of the network (they are “orthogonal”), proving they encode significantly different image features. We also see that StarDist displays considerably less orthogonal deep layers than Cellpose and is altogether more homogeneous. With corruptions, the similarity of shallow blocks is mostly preserved. The inter-layer similarity structure of StarDist is mostly unchanged Geometric and Digital corruption types, and more contrasted for Noise, Blur and Assays. On the other hand, this structure collapses for Cellpose in Assays (all layers encode the same information) and the information is saturated in the early layers of Blur and Noise (later layers encode more or less the same information), destroying the deeper layers’ differentiation capacity. The deeper layers’ orthogonality is better preserved for Digital and Geometric corruptions.

Overall, this figure shows that the layers of the less robust neural network lose their ability to encode complementary information in the presence of corruption. We also posit that the shortness of StarDist’s neural network might also be a contributing factor to its robustness: since it contains a lesser number of orthogonal intermediate layers than Cellpose, the propagated information is more similar throughout the layers, and losing one portion of the information bears a lighter prejudice to the overall performance; whereas Cellpose might be more reliant on these complementary (but fragile) orthogonal encodings. Also, the stronger regularity prior of StarDist (namely that it favors star-convex segmentations) probably strengthens intra-layer similarities and, therefore, its overall robustness.

We assess in figure 4b the independent robustness (per corruption type and severity level) of 5 layers of different depths: two early layers (10th and 30th percentiles) taken from the encoding path, one intermediate layer (50th percentile) from the bottleneck, and two later layers (70th and 90th percentiles) from the decoding path. This figure reports the CKA values of each layer to itself, computed on clean and corrupted images: in other words, it measures how much a layer is similar to itself in the presence of different corruption types and severity levels (intra-layer similarity). Overall, the intra-layer similarity of each layer decreases with severity level. For Cellpose, the intra-layer similarity of the 10th percentile layer declines with varying speed according to the corruption type: it does so strongly with Noise, slowly with Geometric, and mildly with Blur, Digital and Assays. StarDist features the same patterns, although with greater robustness: the intra-layer similarity if almost always optimal for Geometric and Digital, declines significantly more slowly for Noise and Blur, and is comparable in Assays. The difference in the other layers’ robustness if less pronounced, suggesting a particular importance of the early steps of the encoding stage in the overall robustness of a segmentation network.

We visually confirm this intuition in Figure 4c, where we illustrate the feature maps produced by each of these layers for both models, on a fluorescence microscopy image taken from the BBBC019 dataset. Here, we use a PCA projection over the three largest components of the channel axis to turn each feature map into an RGB image. Both methods show good performance on the clean version of this image. Yet, we see that the intermediate representations are much less stable in the case of Cellpose. For Cellpose, the early 10th percentile layer and the later 70th and 90th percentile layers look like pure noise with Noise corruption. Cellpose is able to retrieve cell-related features with Blur corruptions, but the boundaries are irretrievably mingled. As for Digital corruptions, only boundary-related features are encoded in its feature maps. StarDist’s feature maps are strikingly different. Early and later layers are all visually interpretable across all corruption types. Contrary to Cellpose, cell-like features are spotted across all layers for Noise, they can be clearly delineated for Blur, and the interior of the cell is likewise encoded for Digital. Only the intermediate layer (50th), which seems to encode background locations in the clean image, is deficient in robustness and is cleanly retrieved only for Digital.

## Discussion

Our benchmark for biological segmentation models establishes three principal findings whose implications extend beyond the specific models evaluated.

1. performance on clean images is substantially decoupled from robustness to corruptions. This is in contrast to the results of Kamann and Rother (12) for semantic segmentation on natural images, but aligns qualitatively with Hendrycks and Dietterich (11)’s original classification finding on natural images. The difference likely reflects the highly heterogeneous semantic properties of cell imaging compared to natural images and the distinct nature of the instance-level cell detection task, which is simultaneously sensitive to photometric and geometric corruptions in greater ways than semantic segmentation and class-level predictions are.
2. StarDist displays higher robustness to image corruptions than Cellpose and most SAM-based competitors, despite inferior overall F1-score performance. This result is counterintuitive from the perspective of the scaling hypothesis — the idea that more training data and larger architectures should yield better-calibrated models with more robust predictions. We attribute StarDist’s advantage to the strong inductive bias incurred by its geometric prior: by constraining cell representations to star-convex polygons, the model is forced to learn feature detectors that are selective for the shape and boundary of individual cells rather than for photometric patterns that will be lost with image corruption. This hypothesis is supported by our layer-wise CKA analysis, which shows that StarDist’s representations are substantially stable to corrupted inputs across layers of the encoding path. It is also consistent with the analogous finding of Hendrycks et al. (13) that self-supervised pretraining — another form of inductive bias — provides robustness benefits that do not translate into proportional accuracy gains.
3. training augmentation with a narrow set of corruption types does not confer general robustness: Cellpose-SAM’s inclusion of five augmentation types during training produced robustness gains limited to the matching corruption categories (Noise), but not for the full breadth of Blur or Assay corruptions tested in our benchmark. This suggests a fundamental limitation of augmentation-based robustness strategies when applied to unseen heterogeneous corruption distributions, and echoes the finding of Hendrycks and Dietterich (11) that augmentation with specific corruptions yields fragile, corruption-specific robustness. A more principled approach, such as adversarial training over a broad corruption set, regularisation, or explicit robustness constraints on intermediate representations, may be needed to achieve general robustness.

Our study has several limitations. Our evaluation is restricted to 2D segmentation models and 2D images; the extension to volumetric (3D) microscopy, where corruption patterns differ qualitatively, is an important direction for future work. Our corruption simulations, while physically motivated, necessarily involve parameter choices that may not perfectly match the joint distribution of corruptions encountered in any specific laboratory workflow. In particular, our corruption model assumes spatial invariance and independence between corruption types, whereas real images often exhibit spatially-variable corruptions and corruption interactions. Furthermore, our robustness metrics are averaged over corruption types and severity levels using equal weights; a weighted scheme reflecting the empirical prevalence of different corruptions in specific biological imaging contexts could yield more actionable practical conclusions. Finally, a single corruption type was applied at once, and the study of the robustness of segmentation models to combined corruptions remains to be done.

Our results have concrete implications for the development of future cell segmentation models. The identification of early-layer representational collapse as the primary mechanistic substrate of corruption vulnerability suggests that robustness could be improved by targeted interventions at the feature extraction stage to reduce sensitivity to global photometric statistics — for instance through frequency-domain data augmentation, wavelet-domain normalisation layers, or adaptive instance normalisation strategies. More broadly, the finding that geometric priors confer robustness points to the potential of incorporating other forms of structural domain knowledge — such as size distributions, shape compactness constraints, or topological persistence regularisers — into the training objectives of next-generation foundation models. Our benchmark results and our release of the associated codes provide a principled resource for evaluating such improvements.

## Methods

Our protocol for sampling images, simulating corruption and evaluating segmentation models was done on Python 3 (22), with the pytorch (23), numpy (24), scipy (25), scikit-image (26), opencv (27), tifffile (28), tqdm (29), pandas (30), csbdeep (31), tensorflow 2 (32) packages. The figures were generated using matplotlib (33) and jupyter-notebook (34).

### Dataset curation

#### Source datasets

We sampled images from 30 public biological segmentation benchmark datasets: BBBC039v1 (35), BCCD (36), Cellpose (17), CoNIC (37), CPM15 (38), CPM17 (38), CryoNuSeg (39), DSB2018 (40), DeepBacs (41), DIC-C2DH-HeLa (42), DynamicNuclearNet (43), Fluo-C2DL-MSC (42), Fluo-N2DH-GOWT1 (42), Fluo-N2DH-SIM+ (42), Fluo-N2DL-HeLa (42), IHC-TMA (44), LIVECell (45), LynSec (46), MoNuSac (47), MoNuSeg (48), NeurIPS 2019 (49), NuInsSeg (50), Omnipose (51), PanNuke (52), PhC-C2DH-U373 (42), PhC-C2DL-PSC (42), S-BSST265 (53), TissueNet (43), TNBC (54), YeaZ (55). These datasets are standard benchmarks in biological segmentation and collectively provide the training data for many of the evaluated segmentation models (see Extended Table 2 for cross-references). For fairness of comparison, we sample exclusively from the designated test partition of each dataset when one is available; in the absence of such partition, or in the absence of ground truth segmentation masks in the test set, images are drawn from the training dataset.

#### Imaging modalities

The 30 datasets collectively span 7 distinct imaging modalities:

(1)Brightfield microscopy - present in BCCD, Cellpose, CoNIC, DSB2018, DeepBacs, LIVECell, NeurIPS, Omnipose, and YeaZ; (2) Fluorescence microscopy - present in BBBC019v1, Cellpose, DSB2018, DeepBacs, DynamicNuclearNet, Fluo-C2DL-MSC, Fluo-N2DH-GOWT1, Fluo-N2DH-SIM+, Fluo-N2DL-HeLa, IHC-TMA, MoNuSac, MoNuSeg, NeurIPS, NuInsSeg, Omnipose, PanNuke, SBSS-T265, TissueNet, and TNBC; (3) Phase contrast microscopy - present in Cellpose, NeurIPS, Omnipose, PhC-C2DH-U373, and PhC-C2DL-PSC; (4) Confocal microscopy - present in Cellpose and DeepBacs;(5) Differential Interference Contrast (DIC) microscopy is represented by DIC-C2DH-HeLa, DeepBacs, and NeurIPS; (6) Histological whole-slide imaging (WSI)- present in CoNIC, CPM15, CPM17, CryoNuSeg, DSB2018, IHC-TMA, LynSec, MoNuSac, MoNuSeg, NeurIPS, NuInsSeg, PanNuke, and TNBC; (7) Super-resolution microscopy - present in DeepBacs.

#### Staining protocols

Staining is used in histological and fluorescence imaging. 12 staining protocols are represented. The actual number is certainly higher, as several datasets do not comprehensibly report the stains they use. For histological imaging: Hematoxylin and eosin (H&E) staining covers the full set of images available in CoNIC, CPM15, CPM17, CryoNuSeg, IHC-TMA, LynSec, MoNuSac, MoNuSeg, NeurIPS, NuInsSeg, PanNuke, and TNBC. For fluorescence imaging: DAPI-labelling for stained nuclei is present in Cellpose, DSB2018, DeepBacs, DynamicNuclearNet, Fluo-C2DL-MSC (YFP-TIA-1), Fluo-N2DH-GOWT1 (GFP), Fluo-N2DH-SIM+, Fluo-N2DL-HeLa (H2b-GFP), Omni-pose and S-BSST25. Hoechst labelling is present in BBBC019v1, DSB2018, and Omnipose. FITC (fluorescein isothiocyanate) is present in Cellpose and Omnipose. Additional fluorescent dyes include FtsZ-GFP and Nile Red present in DeepBacs. Haematological staining (Giemsa/Wright-Giemsa and Wright’s stain) is present in BCCD. Cytoskeletal markers (F-actin, *β*-tubulin) are present in DSB2018, YeaZ, DeepBacs, and Omnipose, and multiplex immunohistochemistry staining (CD3, CD20, CD38, CDK4, Cyclin-D1, Ki67, and P53) in IHC-TMA.

#### Cell types

27 distinct biological object categories are covered. Cell nuclei segmentation is targeted by a majority of the datasets from a variety of tissue and cancer types (BBBC019v1, CoNIC, CPM15, CPM17, CryoNuSeg, DSB2018, DynamicNuclearNet, Fluo-C2DL-MSC, Fluo-N2DH-GOWT1, Fluo-N2DH-SIM+, Fluo-N2DL-HeLa, IHC-TMA, LIVECell, LynSec, MoNuSac, MoNuSeg, NeurIPS, NuInsSeg, PanNuke, SBSS-T265, TissueNet, TNBC). Whole cell segmentation is targeted for: HeLa cells (DIC-C2DH-HeLa, Fluo-N2DL-HeLa); mesenchymal stem cells (Fluo-C2DL-MSC); GOWT1 cells (Fluo-N2DH-GOWT1); U373 glioblastoma cells (PhC-C2DH-U373, CTC); pluripotent stem cells (PhC-C2DL-PSC);U2OS cells (BBBC039v1, Cellpose); NIH3T3 cells (Cellpose, DynamicNuclearNet, CTC); and a range of other cultured and cancer cell lines, including HEK293, RAW 264.7, PC-3, and HL60 (DynamicNuclearNet), as well as A172, BT474, BV-2, Huh7, MCF7, SH-SY5Y, SkBr3, and SK-OV-3 cells (LIVECell). Additional targeted biological structures include: white blood cells, red blood cells, and platelets (BCCD); bacteria, including named species such as Staphylococcus aureus, Escherichia coli, and Bacillus subtilis (DeepBacs, NeurIPS, Omnipose); yeast (YeaZ); nematodes (C. elegans, Omnipose) and plant cells (Arabidopsis thaliana, Omnipose); bone-marrow derived macrophages from C57BL/6 mice, pancreatic stem cells, pancreatic exocrine and endocrine cells, and non-cell objects (Cellpose); and neurons (Cellpose, NeurIPS). Specific to histological images, several datasets distinguish finer cell-type taxonomies beyond generic nuclei: CoNIC labels epithelial cells, immune cell subtypes (lymphocyte, plasma cell, eosinophil, neutrophil), and connective tissue cells; LynSec targets B-cell lymphoma cells; TNBC labels normal epithelial, inflammatory, fibroblast, macrophage, adipocyte, invasive carcinoma, and myoepithelial cells; and S-BSST25 targets osterosarcoma cells. Tissue-level segmentation targets additionally include multi-organ tissue sections spanning a broad range of cancer and normal tissue types — including breast, kidney, prostate, bladder, colon, lung, and brain tissue (MoNuSeg, MoNuSac), ten organ types in CryoNuSeg, and over 20 human and mouse organs in NuInsSeg.

#### Number of channels

The images contain a varying number of channels, depending on the number of used stains or the acquisition protocol. We systematically convert all images to a 3-channel RGB-like image, using the following approach: Grayscale images are duplicated across the 3 channels ; two channel images are padded with a zero blue channel ; images with more than 3 channels are converted to grayscale by taking the mean channel-wise; and then extended to 3 channels via replication.

### Image sampling strategy

Selecting a maximally representative subset of images from the 30 datasets is a non-trivial optimization problem. The most common strategy, a simple random sample, runs the risk of missing under-represented data (DSB2018 for example is severely imbalanced) as well as over-sampling datasets that contain a lot of redundant images (Dynamic Nuclear Net for example includes frames of videos of slowly evolving cells). We propose a systematic image sampling strategy that optimizes the diversity of the sampled images, via a two-step greedy farthest-point sampling (FPS) in the embedding space of a pre-trained vision foundation model.

All images are first encoded using the ViT-S/14 architecture of DINOv2 (16), a transformer-based model pretrained via self-supervised distillation on a total of 142M natural images. Its encoding capacity in a 384-dimensional latent space has been shown to capture both low and high level semantic image properties. In the first sampling step, we apply greedy FPS independently within each dataset to select 10 images per dataset. FPS is initialized with a randomly-chosen image, and then iteratively selects the most dissimilar image compared to all the previously selected ones, measured as the maximum cosine similarity to the current selection set:

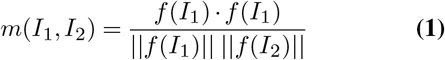

given two images *I*_1_, *I*_2_ and the DINOv2 encoder denoted *f*. We exclude images that do not count at least 5 ROIs in their ground truth. This first step prevents larger datasets

for being oversampled and ensures that the 10 sampled images per dataset span the widest possible range of the dataset’s internal semantic variability (see Supplementary Figure S2). In the second step, FPS is applied globally across all datasets simultaneously, using the 10 per-dataset seeds as initialization points, to select a target final number of images (here 500) from the joint pool. This second step ensures that datasets with greater heterogeneity are sampled more frequently. As shown in Figure 1c, our set of 500 sampled images exhibit a broad and balanced coverage of the full image diversity present in the pool of benchmark cohorts.

### Corruption simulation

We simulate 36 corruption types across 5 physically-motivated categories (see Supplementary Figure S1). Each corruption is parameterised by empirically tuned internal parameters to generate 5 distinct severity levels *s* ∈ {1, 2, 3, 4, 5,} defined as follows: severity 1 is no obstacle to a trained observer, severity 3 represents a level of degradation commonly encountered in suboptimal but scientifically acceptable acquisitions, and severity 5 deliberately exceeds standard quality thresholds in order to test failure modes (see Supplementary Figures S4-S8). The parameter values for each corruption type and severity level are reported in Supplementary Table 1. Corruption types that rely on the cell segmentation mask (e.g. background, photobleaching) receive the ground-truth segmentation mask as an additional input. Geometric corruptions that alter spatial coordinates (barrel, Möbius, mustache, pincushion, shear, stitching, swirl, wavefront) are also applied to the mask using nearest-neighbour interpolation to preserve label integrity. Each image is normalized to the [0, 1] range before turned into a 3-channel RGB-like channel as previously described. We denote the normalized 3-channel image *I*, and its corrupted counterpart *I*^*′*^.

### Noise

The Noise corruption category models stochastic degradations that arise upstream of the analog-to-digital converter, before the acquisition is turned into pixel values.

#### Gaussian noise

arises from thermal fluctuations in the camera sensor, electronic readout noise, and amplifier noise. It is independent of signal intensity and therefore affects equally dim and bright regions. It is modeled as an additive zero-mean normal distribution whose standard deviation is a fraction *σ*_*f*_ of the image dynamic range: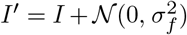The parameter *σ*_*f*_ ranges from 0.1 to 0.5.

#### Shot noise

arises from the discrete, stochastic nature of photon detection. Because photon arrivals follow a Poisson process, the noise variance scales with the local signal intensity. It is simulated as *I*^*′*^ = Poisson(*s* · *I*) */ s*. The scale parameter *s* ranges from 20 to 3.

#### Speckle noise

models multiplicative coherent-light interference, characteristic of laser illumination. The noise amplitude scales proportionally with local intensity, so bright objects acquire granular texture while dark backgrounds remain comparatively clean. It is implemented as *I*^*′*^ = clip(*I* + *I* ·*N* (0, *σ*^2^)), with the clip operator used to preserve the initial image intensity range. The parameter *σ* ranges from 0.3 to 1.5.

#### Salt-and-pepper noise

also known as impulse noise, models transient detector failures, or corrupted pixel readouts, which set individual pixels to the maximum or minimum intensity value. A fraction *p* of pixels is randomly set to either 0 (pepper) or the image maximum 1 (salt), with equal probability. The parameter *p* ranges from 0.025 to 0.125.

#### Hurl noise

generalises impulse noise by replacing a fraction *p* of pixels with values drawn uniformly at random from the image dynamic range, rather than restricting replacements to the two extreme values. This models a broader class of readout malfunctions. The parameter *p* ranges from 0.1 to 0.5.

#### Mixed noise

combines shot noise, Gaussian noise, and a fixed-pattern structured noise (see Pattern noise) component with fixed weights (0.45, 0.45, 0.10). This models real acquisition conditions in which multiple noise sources co-occur, as typical in fluorescence microscopy where Poisson photon noise and Gaussian readout noise are both present and occasionally modulated by camera-specific fixed-pattern artifacts.

### Blur

The Blur corruption category models the loss of spatial resolution arising from imperfect convergence of the optical wavefront at the focal plane. We simulate one standard Gaussian blur and 12 physically-grounded optical aberrations derived from the Zernike polynomial basis of the wavefront.

#### Gaussian blur

provides a standard, spatially isotropic reference blur corresponding to mild defocus or the use of an anti-aliasing filter. It is implemented by convolving the image with a Gaussian kernel of standard deviation *σ* = *σ*_*f*_ · max(*H, W*), where *H* and *W* are the image dimensions. The parameter *σ*_*f*_ ranges from 0.01 to 0.022.

#### Zernike-based optical aberrations

Optical aberrations arise when the wavefront of light converging on the focal plane deviates from a perfect sphere, causing the point spread function (PSF) of the system to become extended and asymmetric. The Zernike polynomials 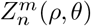 form a complete orthogonal basis on the unit disk, making them the natural representation for wavefront distortions in optical systems (56). Using the concept of normalized pupil coordinates (*ρ, θ*), a Zernike polynomial of radial order *n* and azimuthal frequency *m* is defined as:

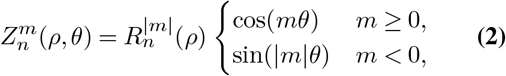

where the radial polynomial is

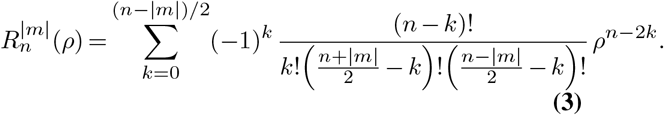

The wavefront aberration *W* (*ρ, θ*) is then expressed as a weighted superposition of Zernike modes,

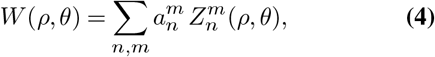

where the coefficients 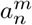 (in units of wavelengths) quantify the contribution of each aberration type. Physically, *n* determines the order of the radial dependence of the aberration (and hence the spatial complexity of the resulting PSF), while *m* encodes the rotational symmetry: *m* = 0 yields rotationally symmetric aberrations (defocus, spherical), |*m* = 1|asymmetric coma-type aberrations, |*m*| = 2 two-fold astigmatic patterns, |*m*| = 3 three-lobed trefoil patterns, and| *m* = 4| four-lobed quadrafoil patterns. Higher radial orders *n* correspond to higher-order variants of the same symmetry class.

To compute the PSF corresponding to a given aberration, we construct the complex pupil function *P* (*ρ, θ*) =]__*ρ≤*1_^·^exp(2*πi W* (*ρ, θ*)*/λ*), where *λ* is the imaging wavelength. The PSF is then obtained as the squared modulus of the Fourier transform of *P* :

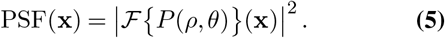

The corrupted image is produced by convolving the original image with the normalised PSF. Our implementation models a circular pupil of diameter 3 mm and a reference wavelength of 450 nm. We simulate the following 12 Zernike modes, identified by their (*n*, | *m*|) indices and standard nomenclature: **defocus** (2, 0), **astigmatism** (2, 2), **coma** (3, 1), **trefoil** (3, 3), **spherical** aberration (4, 0), **second astigmatism** (4, 2), **quadrafoil** (4, 4), **second coma** (5, 1), **second trefoil** (5, 3), **second spherical** (6, 0), **third astigmatism** (6, 2), and **second quadrafoil** (6, 4). Their PSFs are illustrated in Supplementary Figure S3. The severity level for all Zernike-based corruptions is controlled via the aberration amplitude *a* (in *μ*m), scaled proportionally to the image size to maintain consistent visual severity across images of different resolutions: *a* = *a*_*f*_·min(*H, W*)*/*256. The parameter *a*_*f*_ ranges from 0.02 to 0.08 *μ*m.

### Geometric

The Geometric corruption category models spatial distortions of the image arising from a misalignment between the sample and the optical axis, mechanical instabilities, or non-linear optics.

#### Barrel distortion

is a radial distortion in which image magnification decreases with distance from the optical axis, causing straight lines near the edges to bow outward. It is implemented via interpolation of pixel values on a deformed grid using the polynomial radial grid deformation model 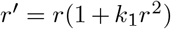 with *k*_1_ *>* 0, where *r* is the normalised radial distance from the image centre. The parameter *k*_1_ ranges from 0.75 to 3.75.

#### Pincushion distortion

is the complement of barrel distortion, in which magnification increases with radial distance, causing straight lines near the edges to bow inward. It uses the same radial model but with *k*_1_ *<* 0. The parameter *k*_1_ ranges from ™0.01 to ™0.20.

#### Mustache distortion

combines barrel and pincushion effects by using both a positive first-order and a negative second-order radial coefficient: 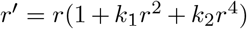 Near the centre the barrel effect dominates; near the edges the pincushion effect takes over, creating the characteristic “moustache” profile. The two coefficients (*k*_1_, *k*_2_) are varied symmetrically. The parameters (*k*_1_, *k*_2_) range from(0.2, ™0.2) to (0.6, ™0.6).

#### Shear distortion

is a linear affine distortion that displaces image rows proportionally to their column position (or vice versa), simulating the effect of a laterally tilted sample or a scanning artefact. It is implemented via the affine transformation matrix 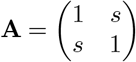with equal shear factor *s* in both directions. The parameter *s* ranges from 0.2 to 0.6.

#### Möbius distortion

is a non-linear conformal mapping of the image plane, defined in complex coordinates *z* = *x* + *iy* as *f* (*z*) = (*az* + *b*)*/*(*cz* + *d*) with *a* = *d* = 1, *b* = 0, and *c* used to control the severity level. Unlike the radial and affine corruptions, Möbius mappings are topology-preserving but can introduce severe local magnification variations and boundary warping, making them substantially harder for segmentation models. The parameter *c* ranges from 0.005 to 0.025.

#### Swirl distortion

applies a spatially decaying rotational displacement in polar coordinates (*r, θ*) whose angular offset decreases with radial distance from the image centre according to 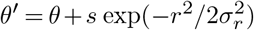, where *σ*_*r*_ = 150 px and *s* controls the severity level. The result is a vortex-like pattern that is most pronounced near the image’s edges and fades towards the center. The parameter *s* ranges from 1 to 3.

#### Wavefront distortion

applies a sinusoidal horizontal displacement of the form *x*^*′*^ = *x* + *A* sin(2*πy/H*), with displacement amplitude *A*. It models lateral sample displacement or wavefront tilt artifacts in scanning systems. The parameter *A* ranges from 30 to 150 pixels.

### Digital

The Digital corruption category models artifacts introduced after analog-to-digital conversion, during data processing, compression, or transfer.

#### Aliasing

arises when an image is acquired or displayed at a resolution lower than the Nyquist frequency, causing high-frequency content to fold back as spurious low-frequency patterns. It is simulated by downsampling the image by a factor *η* using bilinear interpolation, then upsampling back to the original resolution using nearest-neighbour interpolation. The parameter *η* ranges from 1.5 to 7.5.

#### Banding

(intensity quantisation) arises when the number of available intensity levels is reduced, as may occur with low-bit-depth cameras or aggressive integer compression. It is implemented by quantising pixel intensities into *B* equally-spaced bands, the parameter *B* ranging from 8 to 4.

#### Compression

artifacts correspond to the blocky artefacts and ringing patterns characteristic of lossy JPEG compression. We implement this using the standard JPEG codec implemented in OpenCV with the input quality level parameter used to control the severity level. The quality level parameter ranges from 10 (low compression) to 1 (extreme compression).

#### Saturation

models the clipping and remapping of intensity values. It is implemented as histogram equalization of each channel to generate *Î* followed by a global scaling *I*^*′*^ = *γ* · *Î*, with a scaling factor *γ*, and then clipping to the original image dynamic range. This produces colour and contrast distortions that are particularly severe for multi-channel fluorescence images. The parameter *γ* ranges from 1.3 to 2.5

### Assay

The Assay-specific corruption category models artifacts that arise from the biological and experimental conditions of the sample itself, rather than from the imaging optics or digital pipeline.

#### Background texture

models the contribution of out-of-plane fluorescence, scattered light, or autofluorescence from the sample matrix, which superimposes a spatially structured intensity pattern on the background. It is simulated by adding Perlin noise — a smooth, multi-octave-based coherent noise with spatially structured correlations — to the background pixels of the image (defined by the ground-truth mask). The multi-octave Perlin noise is simulated using the pnoise2 function from the noise library (57). The noise amplitude is scaled to *σ*_*f*_ times the standard deviation of the whole image. The parameter *σ*_*f*_ ranges from 1 to 5.

#### Pattern noise

superimposes a fixed-pattern to the image. It is implemented by adding a deterministic spatial checker-board of amplitude proportional by a factor *s*_*p*_ to the image’s dynamic range. The parameter *s*_*p*_ ranges from 0.25 to 1.25.

#### Photobleaching

models the irreversible photochemical degradation of fluorescent dyes upon prolonged illumination, which causes fluorescence intensity to decay as a function of cumulative light exposure. We implement this corruption as an intensity attenuation that is a power-law function of the normalised distance from the foreground ROI mask: *I*^*′*^ = *I* · (*d/d*_max_)^*γ*^, where *d* is the Euclidean distance transform from the foreground ROI mask, and *γ* is an exponent used to control the severity level. Larger *γ* produces a sharper falloff and greater relative darkening of the background. The parameter *γ* ranges from 1 to 2.2.

#### Stitching artifacts

model the geometric and photometric discontinuities that arise at tile boundaries when large fields of view are assembled from multiple overlapping acquisitions. We simulate this by dividing the image into a *n* × *n* grid of non-overlapping patches, randomly shuffling the patches, and reassembling the grid. A higher number of patches produces more numerous and more locally disruptive discontinuities.The parameter *n* ranges from 2 to 16

#### Uneven illumination

models spatial non-uniformity of the illumination field arising from misalignment of the light source, vignetting, or inhomogeneous sample thickness. It is simulated by multiplying the image with a spatially-varying scale map constructed by bicubic interpolation of random scale factors at a set of 5 randomly placed control points. The scale factors at control points are randomly drawn in a range of 1*/r* to *r* which is used to control the severity level and anchored to 1 at the image corners to avoid global intensity collapse. The parameter *r* ranges from 2 to 10. The positions and scale factors (for a given severity level) of the control points are fixed across images for fair comparison.

#### Vibration

models motion blur arising from mechanical vibrations of the sample plate or the objective during image acquisition. It is implemented as a one-dimensional linear motion blur kernel of length *ℓ* pixels applied at a random orientation angle. The parameter *ℓ* ranges from 7 to 37 pixels.

### Segmentation models

#### Segment Anything Model (SAM)

SAM (20) is a promptable image segmentation foundation model trained on a large corpus of natural images. We evaluate three backbone sizes: ViT-T, ViT-B, and ViT-H, using the automatic mask generation mode with default parameters. These serve as unspecialised baselines with no biological domain knowledge.

#### Cellpose-SAM

Cellpose-SAM (9) is a transformer-based segmentation model that couples a SAM encoder with a Cellpose-style flow-based decoder, and was trained on a large, diverse compilation of biological images including BBBC019, BCCD, the Cellpose 3.0 dataset, CoNIC, CPM15, CPM17, CryoNuSeg, DeepBacs, IHC-TMA, LIVECell, LynSec, MoNuSac, MoNuSeg, NeurIPS 2019, NuInsSeg, Omnipose, PanNuke, TissueNet, TNBC, and YeaZ. Importantly, Cellpose-SAM’s training pipeline includes augmentations for five corruption types: shot noise, Gaussian noise, downsampling, Gaussian blur, and an anisotropic blur. All images are provided as single-channel grayscale, normalised to [0, 1].

#### Micro-SAM

Micro-SAM (19) fine-tunes SAM on a curated set of light microscopy datasets: DSB2018, DeepBacs, DIC-C2DH-HeLa, DynamicNuclearNet, Fluo-C2DL-MSC, Fluo-N2DH-GOWT1, Fluo-N2DH-SIM+,\Fluo-N2DL-HeLa, LIVECell, NeurIPS 2019, PanNuke, PhC-C2DH-U373, PhC-C2DL-PSC, and TissueNet. Multi-channel images are converted to grayscale before embedding by averaging channels. We evaluate three backbone sizes `(vit_t_lm, vit_b_lm, vit_l_lm)` using the automatic instance segmentation function from the official repository.

#### CellSAM

CellSAM (18) adapts SAM for cell segmentation using training data from the Cellpose dataset, CPM15, CryoNuSeg, DSB2018, DeepBacs, MoNuSac, NuInsSeg, Omnipose, TNBC, and YeaZ. We use the `cellsam_pipeline` automated segmentation function from the official repository with default parameters.

#### Cellpose (cyto3)

We evaluate the `cyto3` pre-trained model from Cellpose 3.0 (17), trained on the Cellpose cellular and nuclear datasets, TissueNet, LIVECell, Omnipose, YeaZ, and DeepBacs. Cellpose normalises all images such that the average ROI diameter equals 30 pixels; a biologist-provided estimate of the average diameter is used rather than the model’s automatic estimation, which we found to yield substantially inferior results. All images are loaded as grayscale.

#### StarDist

StarDist (14) detects objects as star-convex polygons described by a set of radial distance vectors from each centroid to the object boundary. We use two pre-trained models from the official repository: `2D_versatile_fluo` (trained on DSB2018) and `2D_versatile_he` (trained on MoNuSac and TNBC). The H&E model is applied to images from BCCD, CoNIC, CPM15, CPM17, CryoNuSeg, IHC-TMA, LynSec, MoNuSac, MoNuSeg, NuInsSeg, PanNuke, and TNBC; all other images are processed with the fluorescence model. Before inference, each image is normalised to [0, 1] by subtracting the 1st percentile and dividing by the range between the 1st and 99th percentiles.

### Performance metrics

#### Instance F1 score

We evaluate all models using an instance-level detection F1 score. Each predicted ROI is matched greedily to the ground-truth ROI with the highest intersection over union (IoU); a match is accepted as a True Positive (TP) if the IoU exceeds a threshold *τ* (default *τ* = 0.5, denoted F1@50). Unmatched predicted ROIs count as False Positives (FP); unmatched ground-truth ROIs count as False Negatives (FN). The F1 score is then:

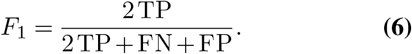

#### Corruption Degradation (CD) and relative Corruption Degradation (rCD)

Adapted from Hendrycks and Dietterich (11), these metrics quantify the degree to which a model’s performance degrades under corruption, relative to a reference model ref (chosen here to be StarDist). Let 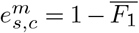 denote the error of a given model *m* at corruption severity *s* and corruption type *c*, where the bar denotes the mean over images. The Corruption Degradation is:

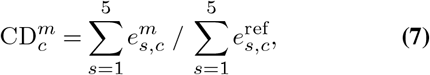

and its mean over all corruption types yields the scalar CD_*m*_. The relative Corruption Degradation corrects for baseline clean-image error *e*_clean_:

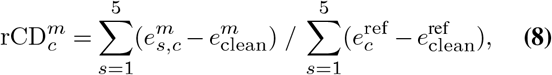

with the scalar rCD^*m*^ defined as its mean over all corruption types. A value of 1 for both metrics corresponds to StarDist performance by construction; values below 1 indicate better-than-reference robustness, values above 1 indicate worse.

### Representation analysis

To analyse how internal representations of segmentation models respond to image corruptions, we extract feature activations from intermediate layers of Cellpose and StarDist for each clean image and each of its corrupted variants. We then measure inter-representation similarity using Centred Kernel Alignment (CKA) (21), a normalised measure of Frobenius-norm similarity between kernel matrices that is invariant to orthogonal transformations and isotropic scaling, making it well-suited for comparing representations of different dimensionalities across layers. For two representation matrices **X** and **Y** (each of shape *n* × *d*, where *n* is the number of images and *d* the dimension of the embedding space), the linear CKA is:

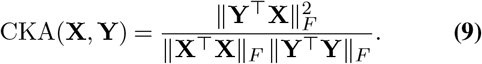

A CKA value of 1 indicates that **X** and **Y** are equivalent up to an orthogonal transformation and rescaling; a value of 0 indicates complete decorrelation. We report CKA between clean representations and corrupted representations as a measure of the representational drift induced by corruption.

## ACKNOWLEDGEMENTS

This work has received funding from the French National Research Agency (ANR) as part of the France 2030 plan (ANR-22-EXES-0013).

