## Supplementary Material for "Benchmarking the robustness of segmentation models to corruptions in biological imaging"

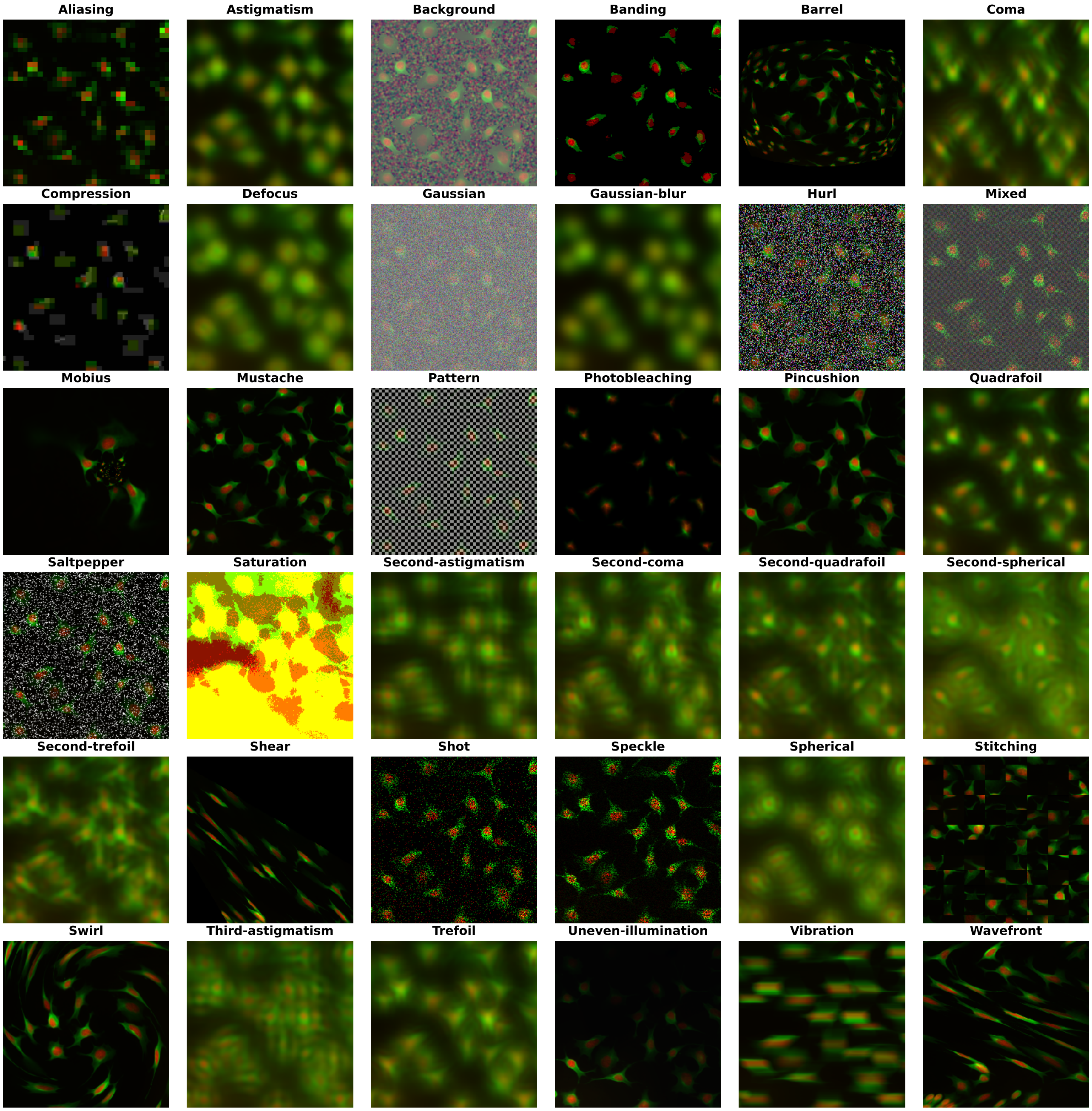


**Fig. S1.** Sample crop of an image from the Cellpose dataset with each corruption applied with a maximum severity of 5.

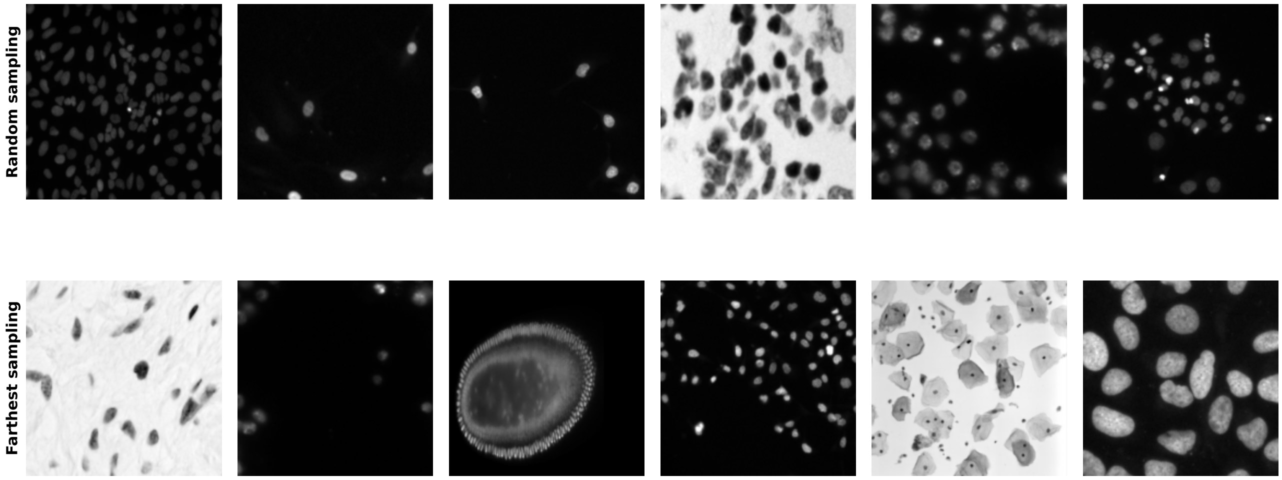


**Fig. S2. Using a basic random sample versus a Farthest Sampling strategy.** The first strategy samples images with similar semantic content (first and fifth column; second and third columns), whilst the other maximizes diversity within a given dataset. Here, all images were taken from the DSB2018 dataset.

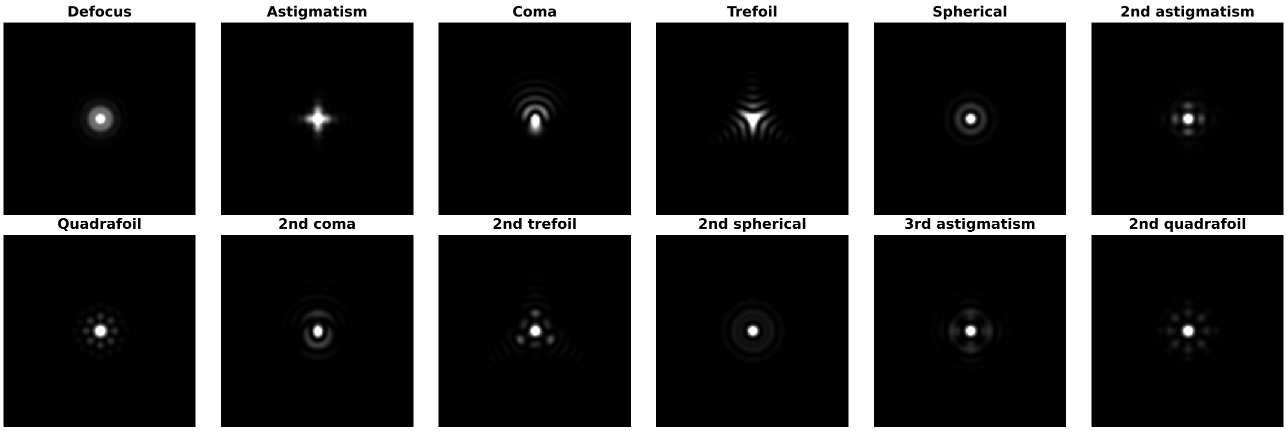


**Fig. S3.** Point Spread Function of the considered Zernike blurs. Kernels were thresholded for better visibility. All kernels correspond to a severity of 5.

**Table S1.** Parameter values for each degradation type across severity levels.

### Digital

| Severity Level | Aliasing | Banding | Compression | Saturation |
|---|---|---|---|---|
| 1 | 1.5 | 8 | 50 | 1.3 |
| 2 | 3 | 7 | 40 | 1.6 |
| 3 | 4.5 | 6 | 30 | 1.9 |
| 4 | 6 | 5 | 20 | 2.2 |
| 5 | 7.5 | 4 | 10 | 2.5 |

### Blur

| Severity Level | Astigmatism | Coma | Defocus | Gaussian blur | Quadrafoil | 2nd astigmatism | 2nd coma | 2nd quadrafoil | 2nd spherical | 2nd trefoil | Spherical | 3rd astigmatism | Trefoil |
|---|---|---|---|---|---|---|---|---|---|---|---|---|---|
| 1 | 0.03 | 0.03 | 0.03 | 0.01 | 0.02 | 0.02 | 0.02 | 0.02 | 0.02 | 0.02 | 0.02 | 0.02 | 0.02 |
| 2 | 0.04 | 0.04 | 0.04 | 0.013 | 0.03 | 0.03 | 0.03 | 0.03 | 0.03 | 0.03 | 0.03 | 0.03 | 0.035 |
| 3 | 0.05 | 0.05 | 0.05 | 0.016 | 0.04 | 0.04 | 0.04 | 0.04 | 0.04 | 0.04 | 0.04 | 0.04 | 0.05 |
| 4 | 0.06 | 0.06 | 0.06 | 0.019 | 0.05 | 0.05 | 0.05 | 0.05 | 0.05 | 0.05 | 0.05 | 0.05 | 0.065 |
| 5 | 0.07 | 0.07 | 0.07 | 0.022 | 0.06 | 0.06 | 0.06 | 0.06 | 0.06 | 0.06 | 0.06 | 0.06 | 0.08 |

### Assays

| Severity Level | Background | Pattern | Photobleaching | Stitching | Uneven illumination | Vibration |
|---|---|---|---|---|---|---|
| 1 | 1 | 0.25 | 1 | 2 | 2 | 7 |
| 2 | 2 | 0.5 | 1.3 | 3 | 3 | 15 |
| 3 | 3 | 0.75 | 1.6 | 5 | 5 | 22 |
| 4 | 4 | 1.0 | 1.9 | 8 | 7 | 30 |
| 5 | 5 | 1.25 | 2.2 | 16 | 10 | 37 |

### Noise

| Severity Level | Gaussian | Hurl | Mixed | Saltpepper | Shot | Speckle |
|---|---|---|---|---|---|---|
| 1 | 0.1 | 0.1 | N/A | 0.025 | 20 | 0.3 |
| 2 | 0.2 | 0.2 | N/A | 0.05 | 15 | 0.6 |
| 3 | 0.3 | 0.3 | N/A | 0.075 | 10 | 0.9 |
| 4 | 0.4 | 0.4 | N/A | 0.1 | 5 | 1.2 |
| 5 | 0.5 | 0.5 | N/A | 0.125 | 3 | 1.5 |

### Geometric

| Severity Level | Barrel | Mobius | Mustache | Pincushion | Shear | Swirl | Wavefront |
|---|---|---|---|---|---|---|---|
| 1 | 0.75 | 0.005 | (0.2, -0.2) | -0.01 | 0.2 | 1 | 30 |
| 2 | 1.5 | 0.01 | (0.3, -0.3) | -0.05 | 0.3 | 1.5 | 60 |
| 3 | 2.25 | 0.015 | (0.4, -0.4) | -0.1 | 0.4 | 2 | 90 |
| 4 | 3 | 0.02 | (0.5, -0.5) | -0.15 | 0.5 | 2.5 | 120 |
| 5 | 3.75 | 0.025 | (0.6, -0.6) | -0.2 | 0.6 | 3 | 150 |

**Table S2.** Training datasets of each segmentation model. The following abbreviations are used: BBBC for BBBC019v1 and CTC (Cell Tracking Challenge) for DIC-C2DH-Hela, Fluo-C2DL-MSC, Fluo-N2DH-GOWT1, Fluo-N2DH-SIM+, Fluo-N2DL-Hela, PHC-C2DH-U373, and PHC-C2DL-PSC.

| Model | BBBC019 | BCCD | CPM15 | CPM17 | Cellpose | CoNIC | CryoNuSeg | CTC | DSB2018 | DeepBacs | DynamicNuclearNet | IHC-TMA | LIVECell | LynSec | MoNuSac | MoNuSeg | NeurIPS | NuInSeg | Omnipose | PanNuke | TNBC | TissueNet | YeaZ |
|---|---|---|---|---|---|---|---|---|---|---|---|---|---|---|---|---|---|---|---|---|---|---|---|
| StarDist | | | | | | | | | ✓ | | | | | | ✓ | | | | | | ✓ | | |
| Cellpose | ✓ | | | | ✓ | | | | | ✓ | | ✓ | | | | ✓ | | | ✓ | | | ✓ | ✓ |
| Cellpose SAM | ✓ | ✓ | ✓ | ✓ | ✓ | ✓ | ✓ | | | ✓ | | ✓ | ✓ | ✓ | ✓ | ✓ | ✓ | ✓ | ✓ | ✓ | ✓ | ✓ | ✓ |
| Micro-SAM | | | | | | | | ✓ | ✓ | ✓ | ✓ | | ✓ | | | | ✓ | | | ✓ | | ✓ | |
| CellSAM | | | | | ✓ | | | | ✓ | ✓ | | | | | | | | | ✓ | | | ✓ | ✓ |
| SAM | | | | | | | | | | | | | | | ✓ | | | | | | | | |

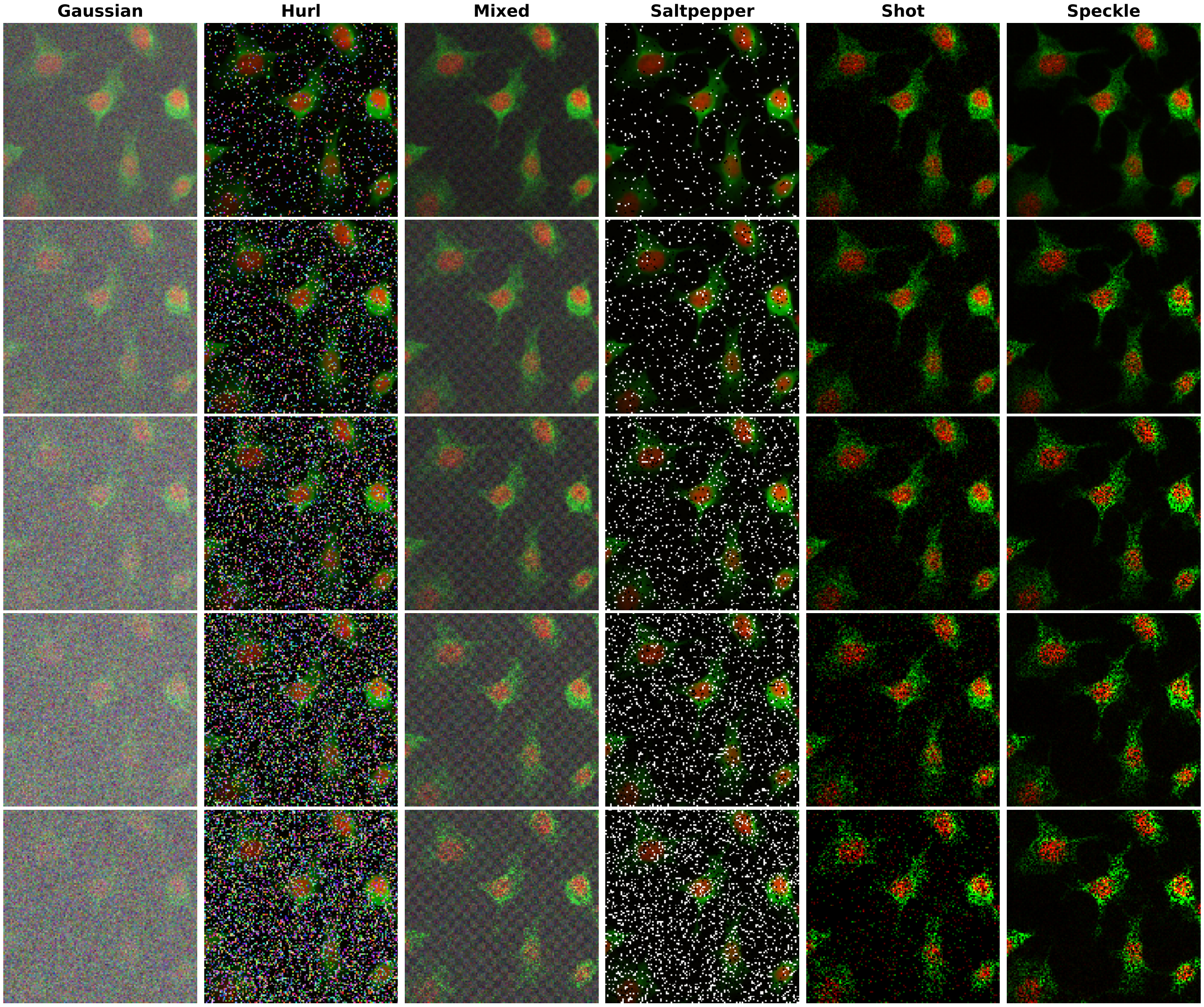


**Fig. S4. Evolution of each Noise corruptions for increasing severity levels.**

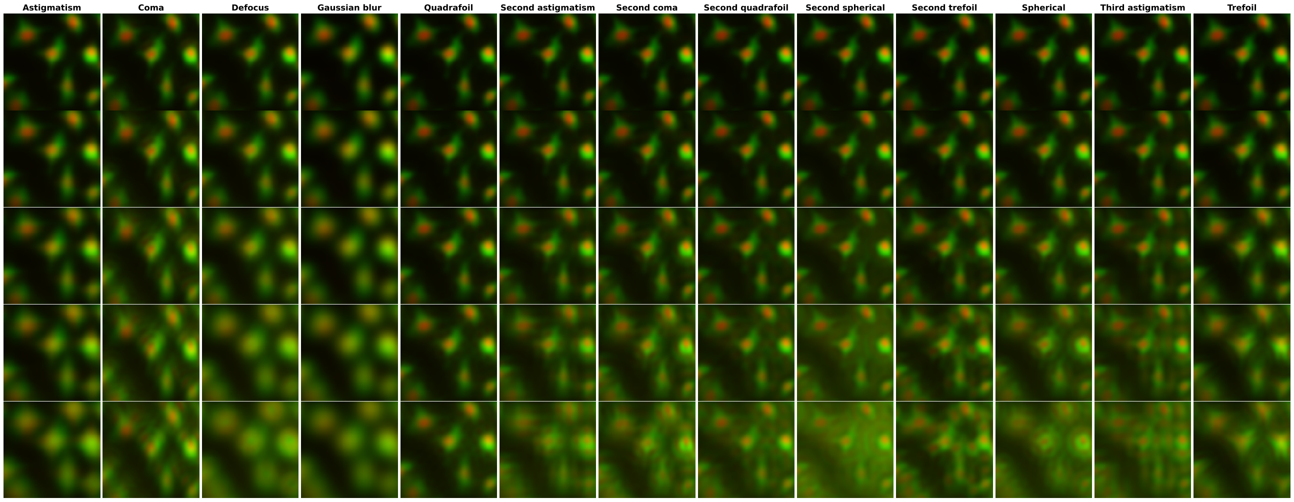


**Fig. S5. Evolution of each Blur corruptions for increasing severity levels.**

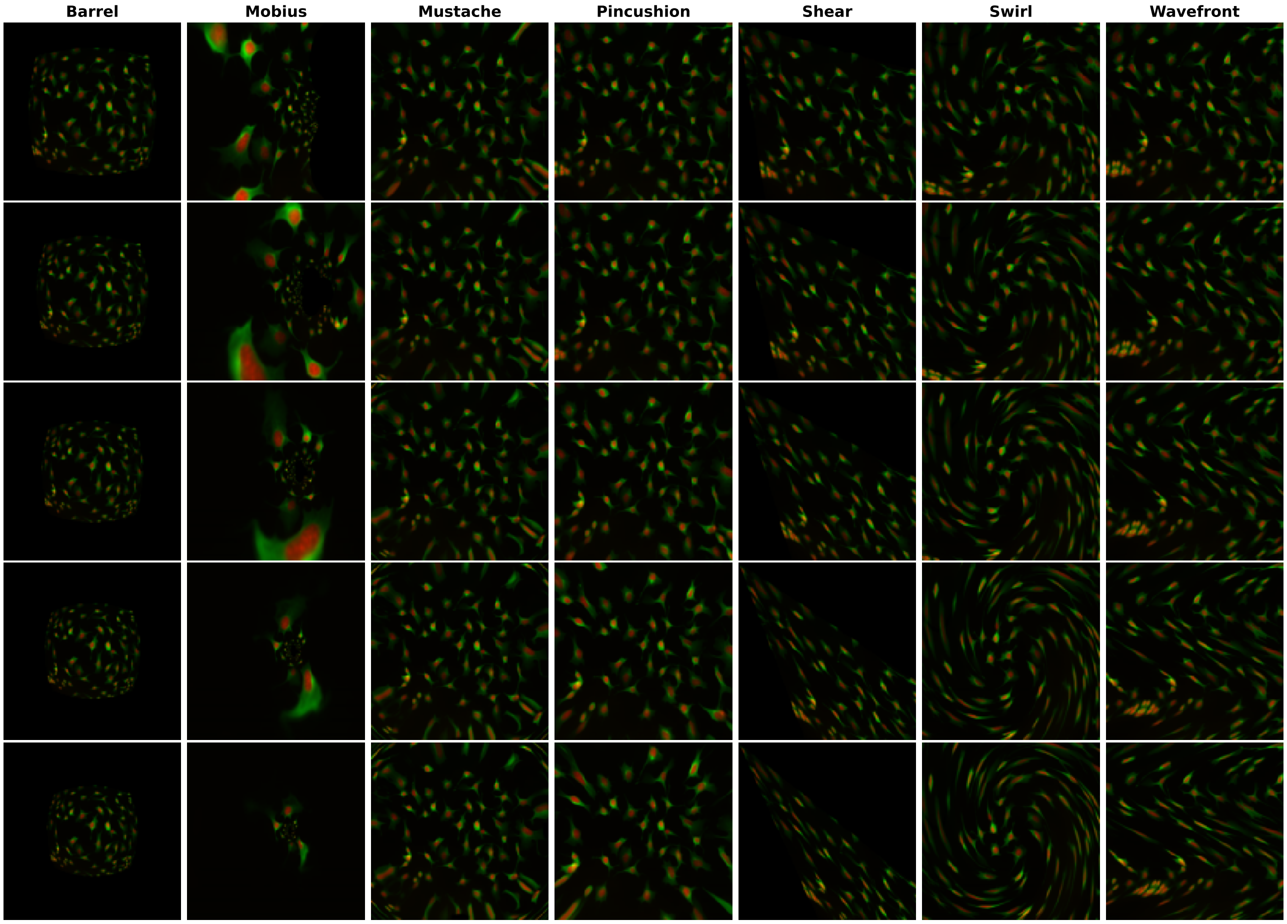


**Fig. S6. Evolution of each Geometric corruptions for increasing severity levels.**

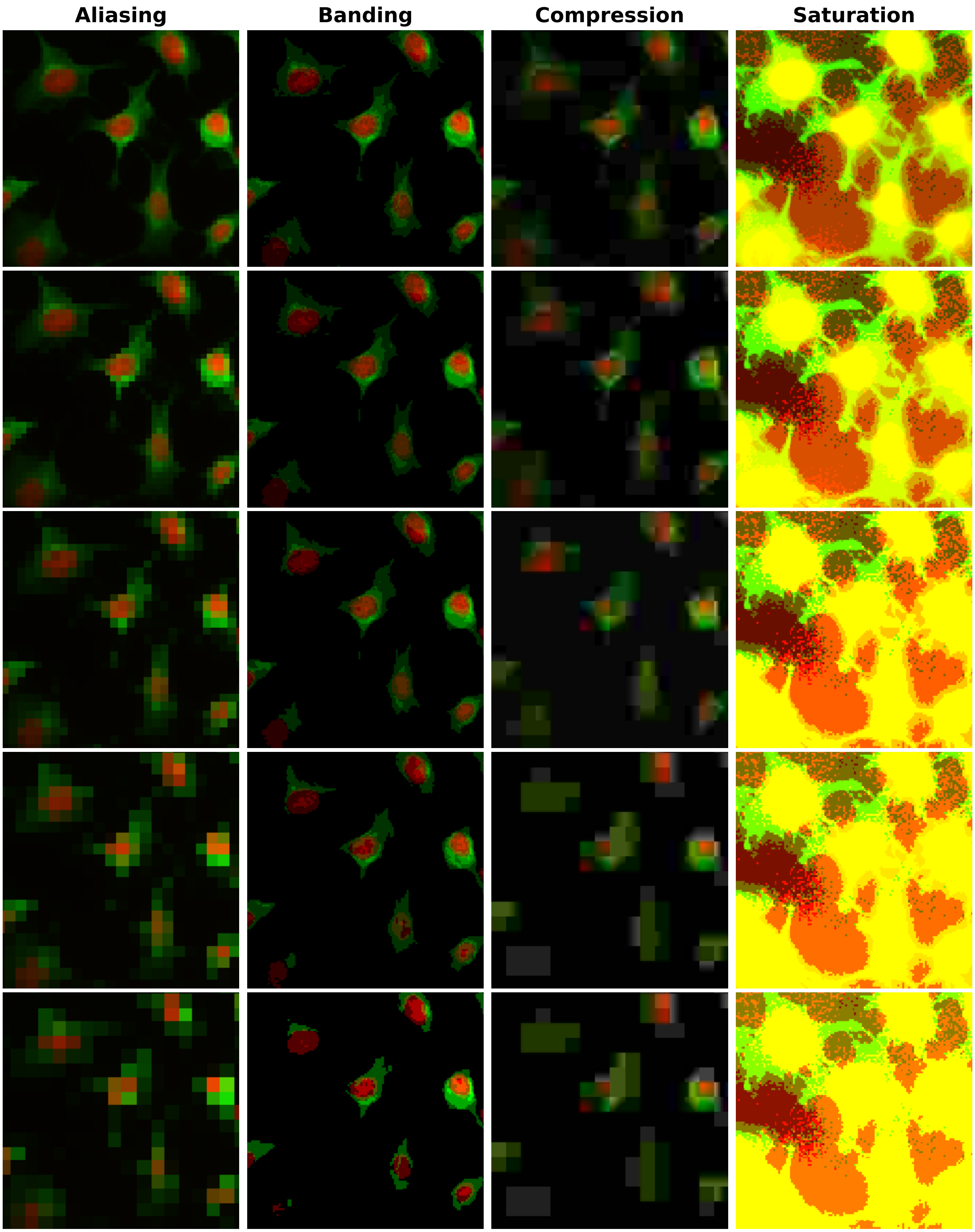


**Fig. S7. Evolution of each Digital corruptions for increasing severity levels.**

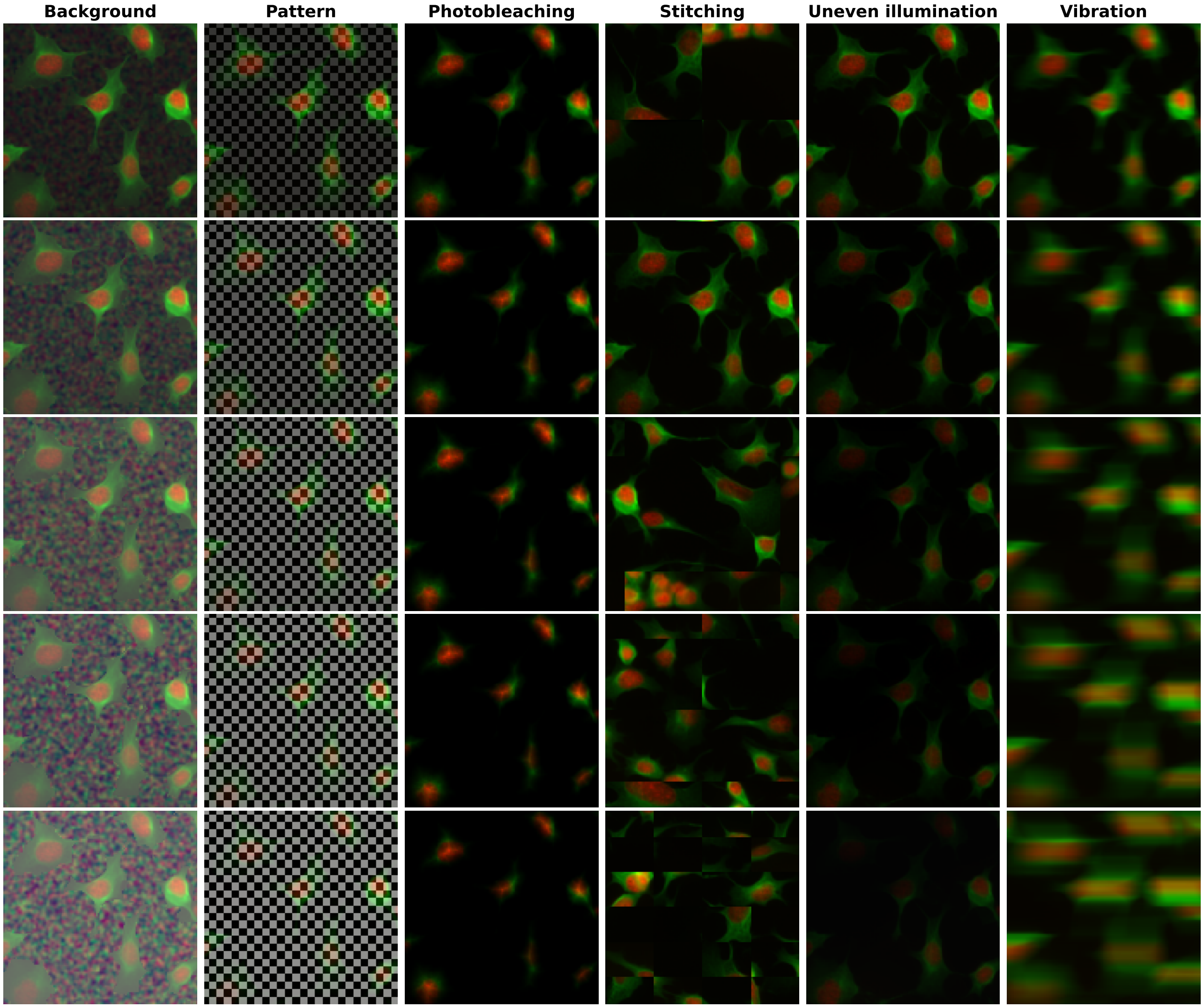


**Fig. S8. Evolution of each Assay corruptions for increasing severity levels.**
